# Nesting Bird Communities of Urban Forest Parks Suffer from Recreational Load (on the Example of Kyiv, Ukraine)

**DOI:** 10.1101/2023.02.10.527978

**Authors:** Tetiana Shupova, Vitaliy Gaychenko, Liudmila Raichuk

## Abstract

The urban environment has a complex effect on the forest diversity. Birds are a suitable object for diagnostic of ecosystem disturbances. The urban recreation load in forest parks was estimated using the author’s methodology according to the total points. To analyze the urban recreation load, we took the characteristics of the forest park and the surrounding landscapes, which are important for the birds’ life, the nesting result, and stable population maintenance: the share of urbanized territory; the share of the territory with unorganized recreation; forest park attendance; the presence of freely rambling pets. The number and distribution of birds were determined by registrations of birds along transects in May–June 2017–2018. 30–54 bird species nest in each forest park with an average density of 1.6 (SD: 0.4)–3.8 (SD: 0.9) pairs/ha. *Parus major* L., *Fringilla coelebs* L., *Turdus merula* L., sometimes *Sturnus vulgaris* L., *Erithacus rubecula* L., *Turdus pilaris* L. are dominant in communities. Urban recreation load leads to a change in the bird nesting strategy, the composition of their communities contributes to the extinction of non-synanthropized species, reduces the abundance of ground-nesters. This leads to a decrease in the number of their populations and some species impoverishment. The synanthropy index of nesting bird communities was 0.38–0.57. A positive correlation between the urban recreation load and the synanthropy index (0.75), Berger–Parker index (0.40), a relative abundance of hollow nesters (0.59), and a negative correlation with the abundance of ground nesters (−0.59) were revealed. To minimize the negative impact of the recreational load on bird communities, to preserve vulnerable species at nesting and maintain the species diversity of avifauna, it is necessary to create areas in forest parks protected from hiking, display in geducational advertising on the importance of preserving animals and their habitats.

## Introduction

In the current century, landscape transformation continues, which affects the stability of the animal habitat (Grimm et al., 2008), leads to the elimination of native species that have not adapted to the impact of the excessive urbanistic press (Ditchkoff et al., 2006), to a reduction in the number of stenotopic bird species, and a decrease in species richness due to the downsizing of some species populations (Martin and Joron, 2003; Felton et al., 2016), and a decrease in species richness due to the downsizing of some species populations. The main threat to songbirds is habitat loss (Lawlor and Meng, 2019).

Taking into account the rather significant transformations of the habitat, perhaps the most significant factors in the survival of avifauna are the possibility of safe nesting and the availability of sufficient food (Runge et al., 2015; Muiruri et al., 2016). It was emphasized that the biodiversity of avifauna is an integral part of ecosystem services, so to ensure the well-being of mankind, global efforts should be focused, amongst other things, on the conservation of bird populations (Wenny et al., 2011).Birds are a particularly beneficial taxonomic group because fulfilling various and important ecological functions in an ecosystem (Fischer et al., 2007; Gardner et al., 2008; Whelan et al., 2008,). To establish the relationships between certain characteristics of the habitat and avifauna transformations, it is especially important to study nesting birds, that is birds at that stage of the life cycle when the connection with the biotope is especially tight and directly depend on all, even small, impacts on the biotope (Blinkova et al., 2020). Changes in the tree species composition can affect the choice of nesting sites by birds (Gabbe et al., 2002), the community structure (Marzluff et al., 1998; Rodewald and Abrams 2002; Blinkova and Shupova, 2017), the overall avifauna diversity (Katsimanis et al., 2006; Heyman, 2010).

In settlements, a significant factor in the formation of nesting bird communities is the change in their natural habitats preserved within the city line (Tomiałojc, 1976; Grimm et al., 2008; Møller et al., 2015). Graham et al. (2014) emphasize that in regions without highly specialized forest bird species, stands of autochthonous tree species can support bird communities comparable to those of natural forests. In the case of simplification and homogenization of the habitat of birds in oak plantations, stenotopic species drop out of the communities (Felton et al., 2016). With insufficient development of the grass cover, which is important as a habitat for invertebrates, insectivorous birds suffer from a lack of food (Batary et al., 2014; Pereira et al., 2014; Bergner et al., 2015). All these features should be taken into account when planning green spaces within the city limits.

The habitat of birds in forest parks is close to the natural one (Paker et al., 2014). But the recreational load makes significant changes in the proper functioning of forest ecosystems, so the violation of forest integrity in Ukrainian cities is a consequence of predominantly urban recreation (Blinkova and Shupova, 2018) as the main anthropogenic factor in this case (Zuniga-Palaciosa Bergner et al., 2020). In addition to the disturbance of the habitat of birds, the impact of human disturbance is also important. It limits the time and space for birds of foraging and breeding, and as a result, stimulates birds to leave highly disturbed fragments (Fernández-Juricic and Jokimäki, 2001; Fernández-Juricic, 2002). Consequently, the need for a multicenter study of bird communities under urban conditions is beyond doubt.

Given that, this study aimed to analyze the changes of both qualitative and quantitative characteristics of nesting bird communities in the forest parks of a megalopolis under different urban conditions to propose measures to mitigate the negative impact of the recreational load on bird communities.

## Material and methods

### Studied Territory

Kyiv (the area is 83560 ha) is located on the line between the Forest and Forest-Steppe Zones on both the right and left banks of the Dnipro River. Whereupon, 31.300 ha ctares of the total area of the city are of natural and semi-natural forests. The northern and north eastern parts of Kyiv are characterized by pine and birch forests. Mixed forest-steppe oak forests are sprawled in the southwest and south of the city. Evaluation of nesting bird communities was carried out on model plots in large forest parks (45–1879 ha), which are objects of the Nature Reserve Fund of Ukraine of national or regional significance (Natural Reserve Fund of Ukraine, 2009). Forest-park plantations are formed of *Quercus robur* L., *Carpinus betulus* L., *Tilia cordata* Mill., *Acer platonoides* L., *Ulmus glabra* Huds., *Pinus sylvestris* L. which dominate the arborescent stratum (Netsvetov et al., 2019). Steppe vegetation, the vegetation of floodplain coastal forests and meadows are represented fragmentarily in forest parks. The valley-gully relief, the presence of a river, a system of streams or lakes are specific for some forest areas. Part of the forest territory has been transformed (deforested, ornamental plants have been planted, a network of paved paths has been made, playgrounds and sports grounds have been equipped). Most of the work on the forest transformation into forest parks was carried out in the 1950-the 60s. In the forests, vacationers spontaneously tread an additional network of paths, there are also areas equipped for picnics, bonfires traces. On the outskirts of the forest areas hospitals, camps for children and youth, sports, and scientific stations are located.

Forest parks are perched inside city limits, surrounded by neighbourhoods of residential or industrial buildings (Fig. 1). Residents of neighbouring microdistricts use them extensively for recreation, as a result of which plots of vegetation regression stages 3 and 4 are registered here (Blinkova and Ivanenko, 2018). In the 1970s and 80s, most forest parks in Kyiv were protected as a biodiversity reserve (Natural Reserve Fund of Ukraine 2009). Some of them attained conservation status in 1990–2009. In 2013, the average recreational load on the national natural parks of Ukraine reached 149 people per day per site or 0.006 people per day per hectare (Zhupanenko, 2014).

**Figure 1.**
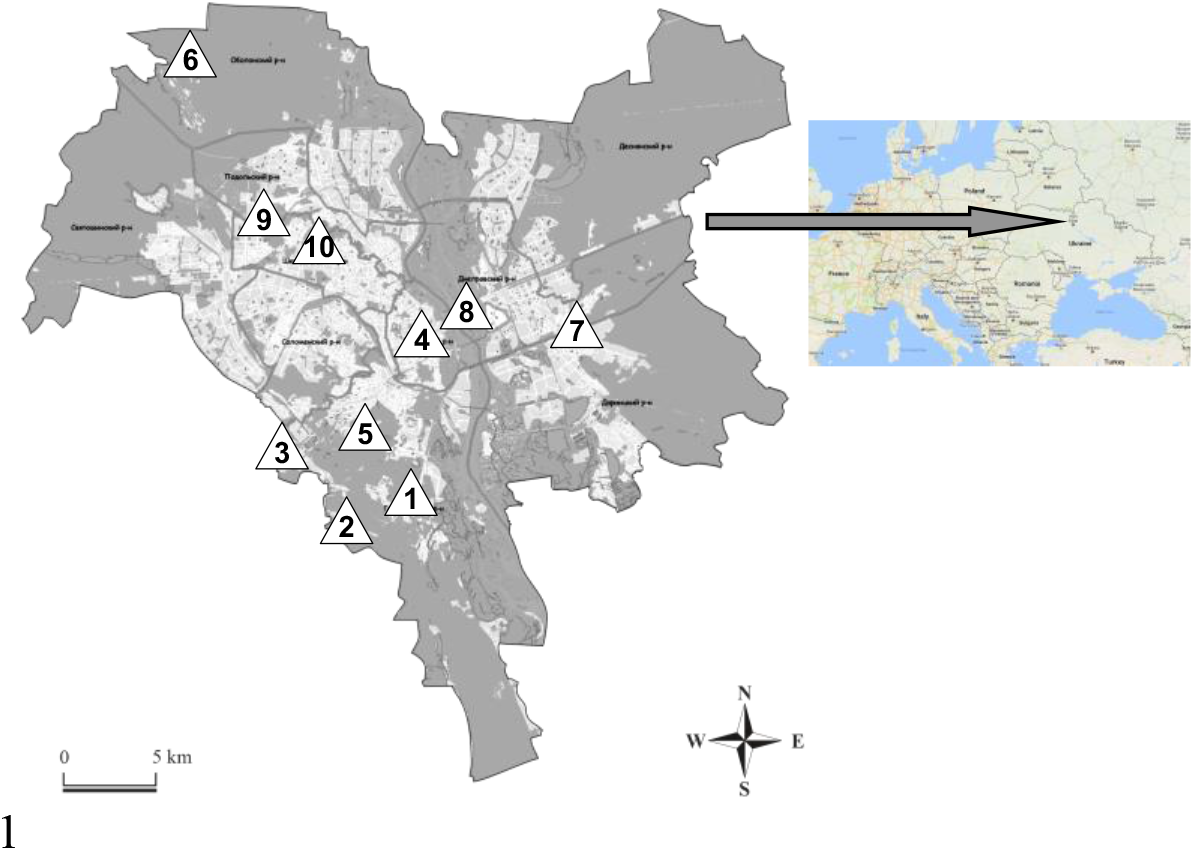
Location of studied forest parks in Kyiv. Note: forest parks: 1 – “Kitaevo”, 2 – “Theophania”, 3 – “Teremky”, 4 – “Lysa Hora”, 5 – “Holosiivskyi Lis”, 6 – “Pushcha Vodytsia”, 7 – “Partisanska Slava”, 8 – “Hydropark”, 9 – “Syretskiy Huy”, 10 – “Nivki”

### Urban Recreation Load Analysis

The urban recreation load in forest parks was estimated using the author’s methodology (Shupova, Koniakin, 2020) according to the total points. To analyze the urban recreation load, we took the characteristics of the forest park and the surrounding landscapes, which are important for the birds’ life, the nesting result, and stable population maintenance:

- the share of urbanized territory: 2 points for every 1–5% of the forest park area under alleys, flower-beds, playgrounds and sports grounds, picnic areas;
- the share of the territory with unorganized recreation: 2 points for every 1-10% of the forest park area with footworn paths, areas of spontaneous picnics, clearings;
forest park attendance: 1 point for 1–5 people/km of transect;
- the presence of freely rambling pets: 1 point for 1 individual/km of the transect; in the neighbourhoods surrounding the forest parks with a radius of about 2 km, the share of the territory of multi-storey buildings was also taken into account. Such biotopes complicate the exchange of individuals between bird communities in neighbouring forest parks. 5 points were given for each 1–10% of the biotopes area that is adjacent to the forest park and occupied by multi-storey buildings.

The following row of forest parks on the gradient of the total urban recreation load points has been compiled: “Kitaevo” → “Theophania” → “Teremky” → “Lysa Hora” → “Holosiivskyi Lis” → “Pushcha Vodytsia” → “Partisanska Slava” → “Hydropark” → “Syretskiy Huy” → “Nivki” (Tab. 1).

**Table 1.** Characteristics of studied Forest parks on the gradient of urban recreation load

| № FP | Name of tract | GPS coordinates | Total points |
| --- | --- | --- | --- |
| 1 | “Kitaevo” | 50°21'48"N, 30°32'26"E | 9 |
| 2 | “Theophania” | 50°20'32"N, 30°29'37"E | 22 |
| 3 | “Teremky” | 50°21'40"N, 30°27'15"E | 36 |
| 4 | “Lysa Hora” | 50°23'42"N, 30°32'53"E | 44 |
| 5 | “Holosiivskyi Lis” | 50°22'01"N, 30°30'30"E | 51 |
| 6 | “Pushcha Vodytsia” | 50°33'05"N, 30°21'10"E | 52 |
| 7 | “Partisanska Slava” | 50°24'44"N, 30°41'01"E | 68 |
| 8 | “Hydropark” | 50°26'27"N, 30°34'44"E | 72 |
| 9 | “Syretskiy Huy” | 50°28'26"N, 30°25'34"E | 76 |
| 10 | “Nivki” | 50°27'40"N, 30°25'13"E | 81 |

### Birds Accounting

For comparison, we selected forest parks, on the territory of which observations were carried out in the same time interval (2017 and 2018). The material was collected in the nesting period (May–June). The number and distribution of birds were determined by the method of transect counts (Bibby et al., 2000) with a transect width of 100 m. The length of the transect was 1.0–1.5 km, and their total length was about 25 km. In small forest parks, the transect length was limited by the length of the forest area. Those forest parks have been surveyed completely. In large forest parks, 2–3 transects were laid depending on the size of the forest park. The registration was carried out annually in May – June in the morning hours (06 a.m. – 11 a.m.). Passeriformes actively were singing. Birds were visible feeding chicks. When planning observations, weather conditions were taken into account: observations were not carried out in the rain and strong winds. If a low activity of birdsong was observed during the registration, the count was repeated 5–10 days later to obtain reliable empirical data. In just 2 years, 38 transect counts were carried out in 10 forest parks. The nesting category included species for which we registered a lekking behaviour, the food collection by adult birds for feeding the chicks, the presence of nests, and the fledgling. For passerines, the presence of a singing male in the studied area was equivalent to the presence of a nesting pair. For each forest park, the average indices of birds’ nesting density for 2 years were struck. The data obtained was used to compare the state of nesting bird communities in ten forest parks. The species of birds and the search for their nests in tree canopies visually were determined using binoculars Nikon Aculon A211/10×50. The acoustic identification of the bird species was also used. The bird species list was presented following the “International Code of the Zoological Nomenclature” (adopted by the International Union of Biological Sciences 2012).

An analysis of the ecological structure of communities was carried out according to Belik (2006). The bird species were divided into different ecological groups depending on their preferred biotopes as habitats. Dendrophils are birds that live in trees and shrubbery plantings. Sclerophiles are species associated with vertically dissected relief: they nest in cracks of soil, stones, stumps, in burrows. The limnophile group includes semiaquatic and waterfowl birds that inhabit a variety of damp habitats (Belik 2006). The nesting strategy was determined by the birds’ choice of nesting microgilds. Dendrophils have various forms of connection with the forest environment: tree canopy nesters (settle in the crowns of tall trees), undergrowth nesters (nest in the crowns of the lower layer of tree-shrubbery plants), tree hollow nesters (in hollows), ground nesters (nest on the ground).The nests of undergrowth nesters are located much lower than those of tree canopy nesters, which nest in the crowns of the main forest-forming tree species. They easily garner people’s attention and are more often ravaged by predators. Therefore, in ecological studies, species that are evolutionarily adapted to undergrowth habitation are distinguished as appurtenant to separate groups of nesting strategy (Shirihai et al., 2001; Camprodon and Brotons, 2006; Belik, 2006).

The number of protected species was evaluated according to the lists of various international conventions, the Red Book of Ukraine, and regionally rare species. The synanthropy index of nesting bird communities was determined according to Jedryczkowski (Klausnitzer, 1990): 

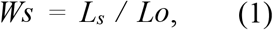

where *L_s_* is the number of synanthropic species, *L_o_*is the total number of species. In this index, we took into account species (populations) that occur in Kyiv city and species (populations) which nest in wild (natural) areas of the Kyiv region.

### Statistical Analysis

The average nesting density of birds was calculated. For average bird density, standard deviation and dispersion were calculated. The standard error and logarithmic trend of analyzed data have been shown in the graphic illustration in Microsoft Excel. Representatives of ecological groups of birds were intercompared by their relative abundance. Several generally accepted ά-diversity indices of communities for each experimental plot were applied (Magurran 1988):

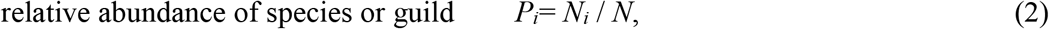

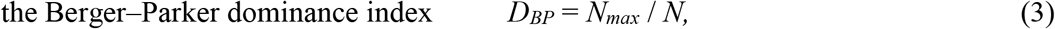

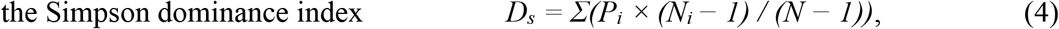

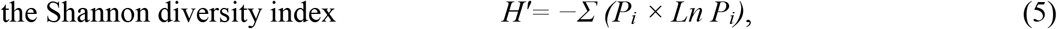

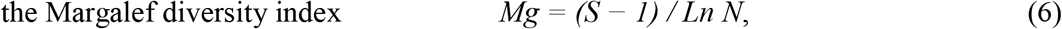

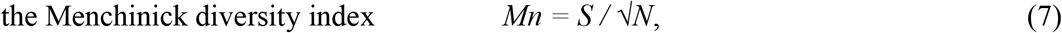

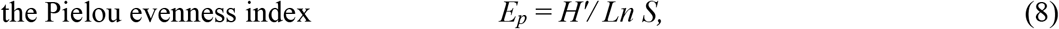

where *N*_i_ – the density of the species (pairs/ha); *N* = *ΣN_i_*– total density; *N_max_* – the maximum value of *N_i_*; *N_m_* – average nesting density; *P_i_* – the ratio of each species; *S* – total number of the species; *S_i_* – frequency of the species; *H’* – Shannon’s index.

Statistical processing was performed in “Origin Pro 9.0”. Principal Component Analysis (PCA) revealed correlations of urban recreation load with data synanthropy index, α-diversity indices, number of the ground nesters and undergrowth nesters, the ratio of hollow nesters and ground nesters. We carried out several cluster analyses to identify groups of similarities in some nesting bird communities of 10 forest parks in Kyiv based on the response of these communities to the urban recreation load. In the beginning, we received the results of 2 cluster analyzes of the similarity of nesting bird communities, carried out on the following data:

1. total number of nesting bird species, the Shannon, Berger–Parker and Pielou indices;
2. a total number of nesting bird species, average bird nesting density, the synanthropization indices of communities, the relative abundance of tree canopy nesters, undergrowth nesters, hollow nesters, and ground nesters.

We have obtained the distribution of 10 bird communities into identical groups with small differences in the similarity distance. Then, we changed the total number of nesting species of the community, on the number of bird species representing each nesting strategy (tree canopy nesters, undergrowth nesters, hollow nesters, and ground nesters) in both matrices for analysis. The results were also very close, but yet slightly different. Therefore, for the final analysis, we combined the following parameters in a matrix: the number of bird species for each nesting strategy: Shannon, Berger–Parker, Pielou, synanthropy indices, average bird nesting density, the relative abundance of tree canopy nesters, undergrowth nesters, hollow nesters, and ground nesters. We believe that the last version of the analysis most accurately reflects the response of bird communities to the urban recreation load in the forest park.

## Results

A total of 76 bird species were observed in 10 forest parks. 70 (93.3%) of these were protected by international conventions (Berne, Bonn, and Washington) and 5 (6.7%) were regionally rare ones. Birds of 67 species (89.3%; n = 75) nest in forest parks, 8 more species from other biotopes visit forest parks for feeding.

From 30 to 54 bird species nest in each of the model forest parks (Tab. 2) with an average density of 1.6(SD: 0.4)–3.8(SD: 0.9) pairs/ha. The logarithmic trend shows a slight decrease in the average bird density in forest parks with an increase in the urban recreation load (Fig. 2). The dispersion of the nesting density of species in communities is high in all forest parks (2.2–6.2), which indicates the presence of urban recreation load.

**Table 2.**
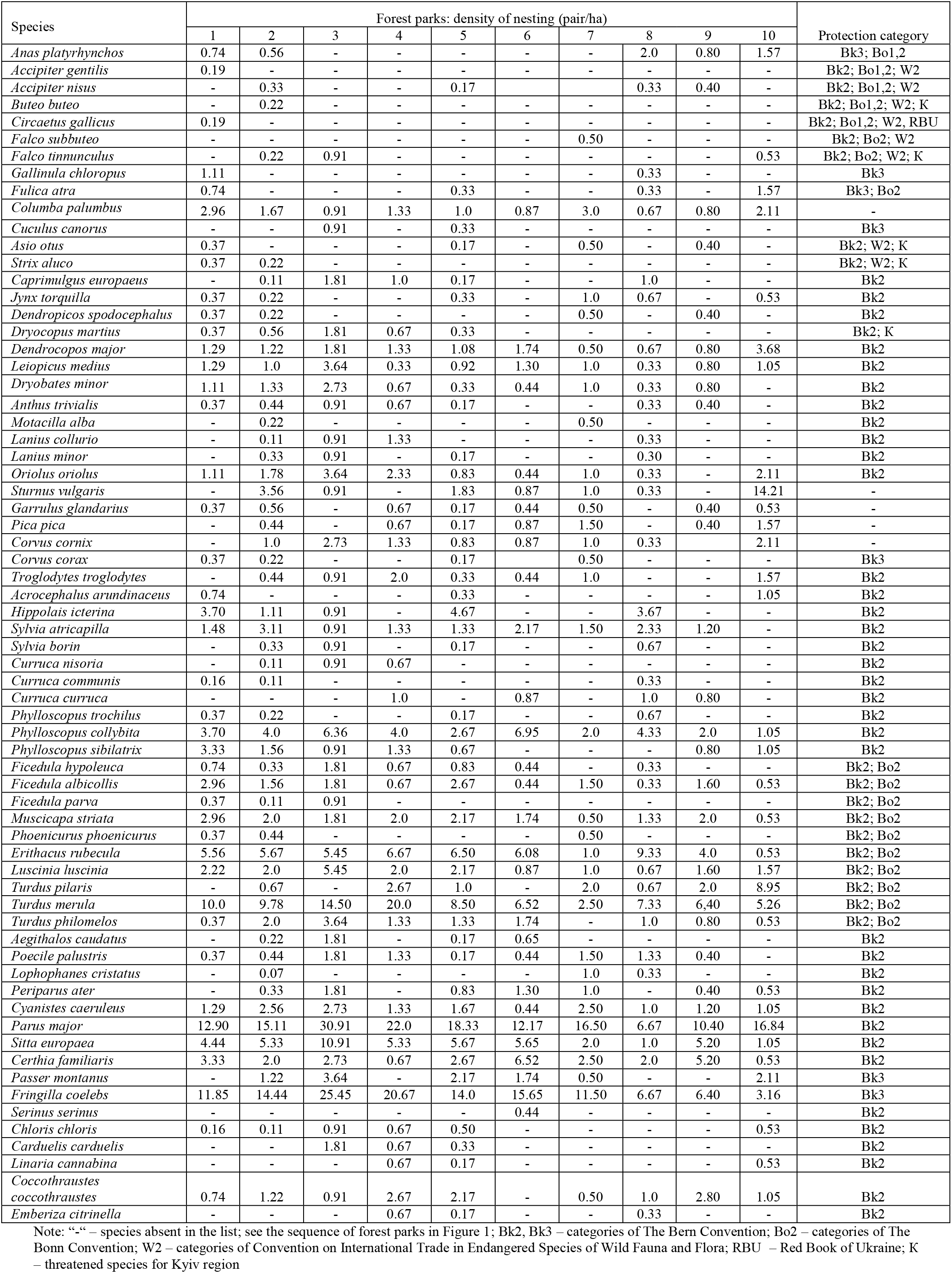
Species composition of bird communities in Forest parks in Kyiv

**Figure 2.**
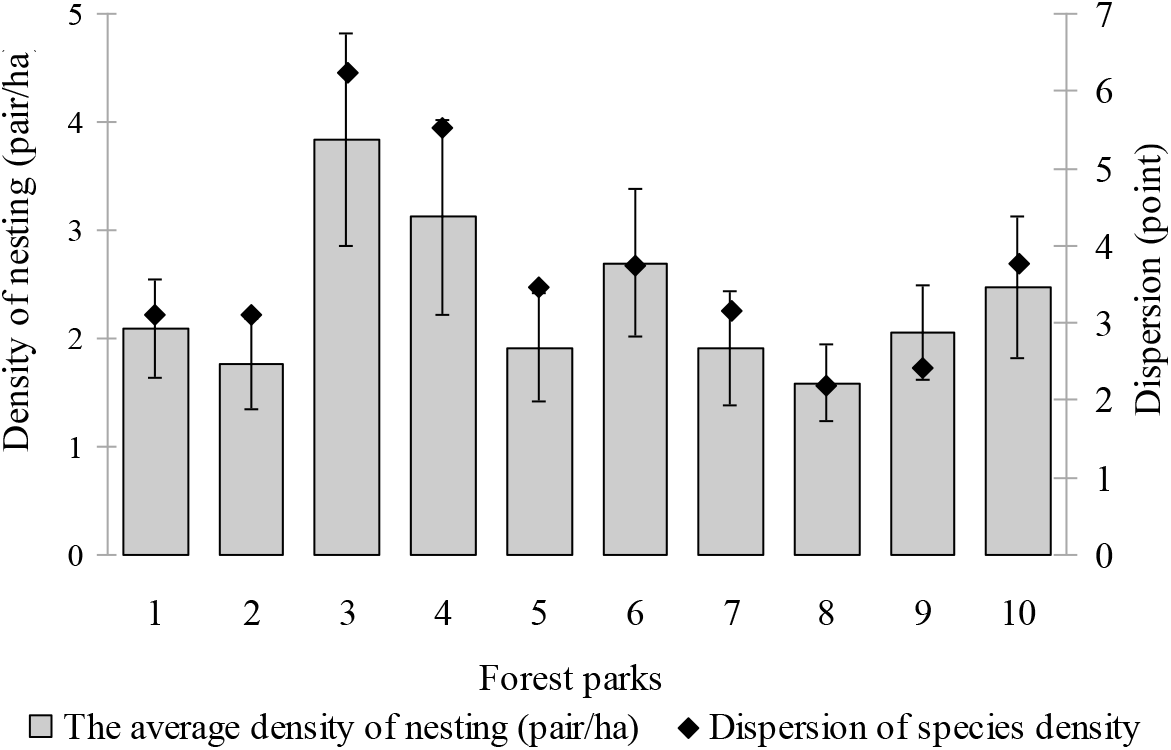
The density of nesting bird communities (SD) and dispersion of the nesting in Forest parks of the Kyiv; see the sequence of forest parks in Figure 1

Blackbird (*Turdus merula* L.), Great tit (*Parus major* L.), Chaffinch (*Fringilla coelebs* L.), Common Starling (*Sturnus vulgaris* L.), European Robin (*Erithacus rubecula* L.), Fieldfare (*Turdus pilaris* L.) dominate almost all forest parks (Tab. 3). As the urban recreation load on forest parks increases the abundance of open canopy nesters (Blackbird and Chaffinch) decreases among the dominants of communities in favour of the cavity nesters, therefore starling and robin become dominants as well as colonynester of field thrush.

**Table 3.** Dominant species in nesting bird communities in Forest parks of the Kyiv

| Species | Relative abundance of species in communities |  |  |  |  |  |  |  |  |  |
| --- | --- | --- | --- | --- | --- | --- | --- | --- | --- | --- |
|  | 1 | 2 | 3 | 4 | 5 | 6 | 7 | 8 | 9 | 10 |
| *S. vulgaris* | 0 | 0.037 | 0.006 | 0 | 0.020 | 0.011 | 0.015 | 0.005 | 0 | **0.174** |
| *Ph. collybita* | 0.042 | 0.042 | 0.040 | 0.035 | 0.028 | 0.086 | 0.030 | 0.069 | 0.032 | 0.013 |
| *E. rubecula* | 0.063 | 0.060 | 0.035 | 0.058 | 0.069 | 0.075 | 0.015 | **0.148** | 0.065 | 0.006 |
| *T. pilaris* | 0 | 0.007 | 0 | 0.053 | 0.011 | 0 | 0.030 | 0.011 | 0.032 | **0.110** |
| *T. merula* | **0.114** | **0.103** | **0.092** | **0.173** | **0.090** | 0.080 | 0.037 | **0.116** | **0.104** | 0.065 |
| *P. major* | **0.147** | **0.159** | **0.197** | **0.191** | **0.195** | **0.150** | **0.246** | **0.106** | **0.169** | **0.206** |
| *S. europaea* | 0.051 | 0.056 | 0.069 | 0.040 | 0.060 | 0.070 | 0.030 | 0.016 | 0.084 | 0.013 |
| *C. familiaris* | 0.038 | 0.021 | 0.017 | 0.006 | 0.028 | 0.080 | 0.037 | 0.032 | 0.084 | 0.006 |
| *F. coelebs* | **0.135** | **0.152** | **0.162** | **0.179** | **0.149** | **0.193** | **0.172** | **0.106** | **0.104** | 0.039 |

Note: dominants of communities are shown in bold; see the sequence of forest parks in Figure 1

Nesting bird communities in forest parks of Kyiv were divided into 3 ecological groups, most of which were dendrophils (81.8–100% of the composition of communities). The proportion of sclerophiles was 0–9.5%, and that of limnophiles – 0–9.1% of the species composition of communities.

Among open nesters, tree canopy nesters, that nest high in the crowns of the main forest-forming tree species, predominate (27.5–36.7%). The share of undergrowth nesters in the species composition of communities is 5.7–18.9%, tree hollow nesters – 32.5–46.7%, and ground nesters – 5.7–16.2% (Fig. 3).

**Figure 3.**
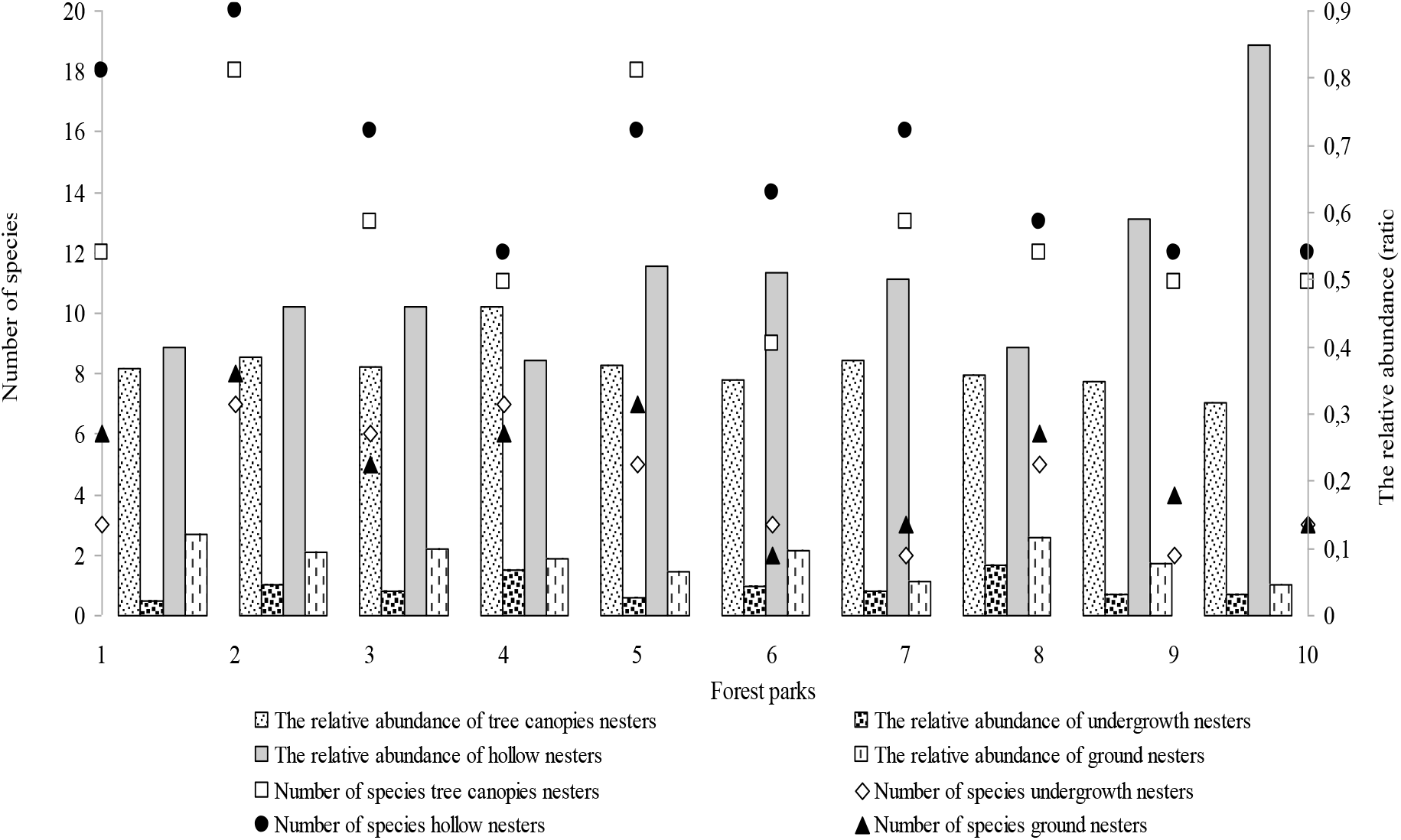
Change in the number of species and abundance of birds of different nesting strategies along the gradient of the urban recreation load; see the sequence of forest parks in Figure 1

The gradient of the synanthropy index of the nesting bird communities of forest parks fits into the parameters 0.38–0.57. The logarithmic trend of relative abundance of synanthropic species shows a slight increase in the gradient of the urban recreation load (Fig. 4). The synanthropy index trend shows a more significant increase.

**Figure 4.**
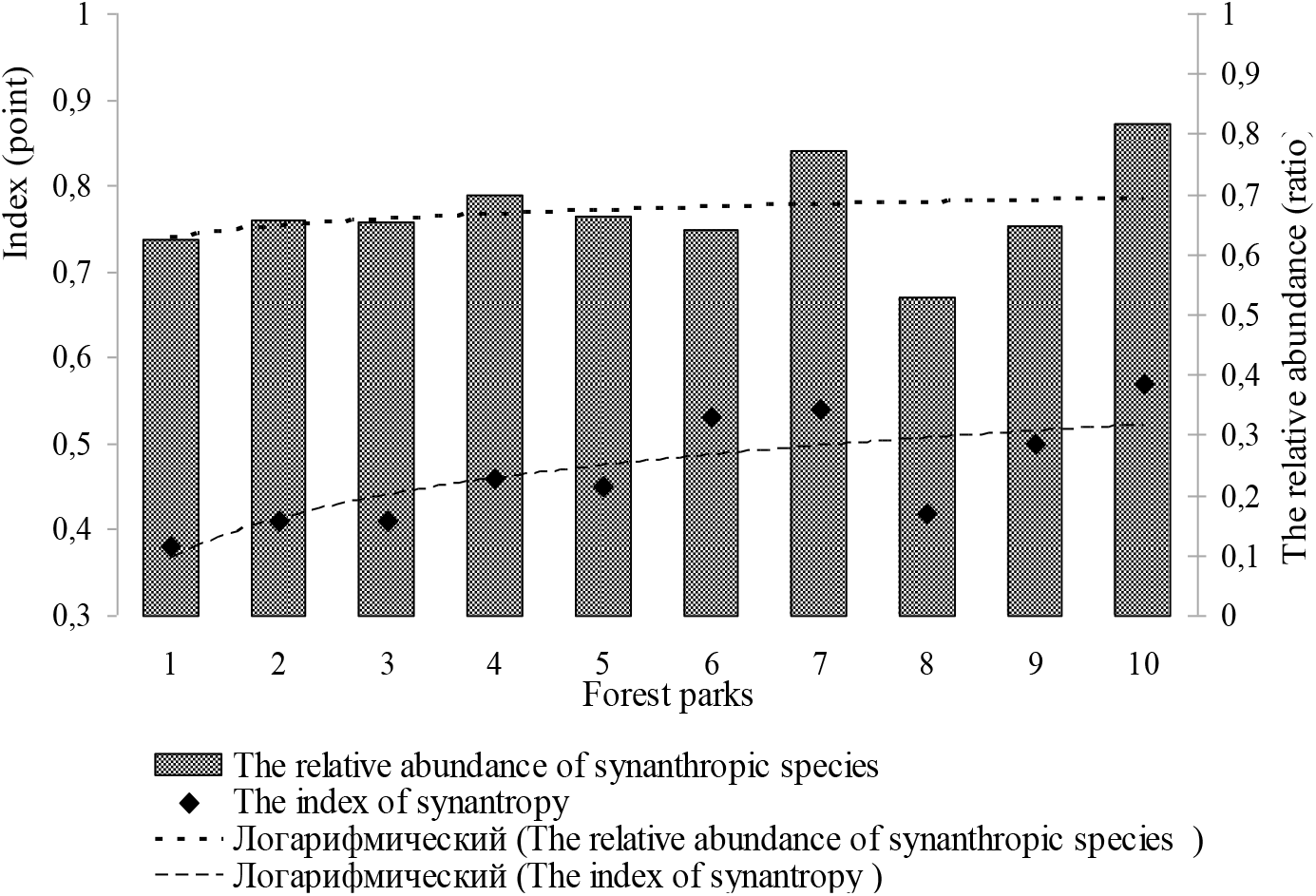
Synanthropization of nesting bird communities in Forest parks of the Kyiv; see the sequence of forest parks in Figure 1

Analysis of the ά-diversity of nesting bird communities showed a rather high species diversity at a low pressure of the dominant species for bird communities of all forest parks of Kyiv. Dominance indices were less variable than diversity indices (Tab. 4). The rates of evennesses of species in communities were high and compatible.

**Table 4.** α-diversity of nesting bird communities in Forest parks of the Kyiv

| Indices | Forest parks |  |  |  |  |  |  |  |  |  |
| --- | --- | --- | --- | --- | --- | --- | --- | --- | --- | --- |
|  | 1 | 2 | 3 | 4 | 5 | 6 | 7 | 8 | 9 | 10 |
| Margalef diversity index | 6.22 | 6.96 | 6.50 | 5.71 | 6.30 | 7.87 | 7.12 | 6.01 | 9.38 | 5.82 |
| Menchinick diversity index | 2.78 | 2.91 | 3.22 | 2.77 | 2.73 | 4.03 | 2.83 | 3.10 | 5.14 | 2.61 |
| Shannon diversity index | 2.82 | 2.91 | 2.90 | 2.65 | 2.98 | 2.95 | 2.88 | 2.78 | 3.10 | 2.71 |
| Simpson dominance index | 0.76 | 0.77 | 0.66 | 0.72 | 0.77 | 0.54 | 0.79 | 0.65 | 0.33 | 0.77 |
| Berger–Parker dominance index | 0.17 | 0.20 | 0.18 | 0.20 | 0.13 | 0.23 | 0.16 | 0.18 | 0.16 | 0.22 |
| Pielou evenness index | 0.81 | 0.80 | 0.85 | 0.80 | 0.86 | 0.83 | 0.79 | 0.84 | 0.82 | 0.80 |

Note: see the sequence of forest parks in Figure

According to cluster analysis, we obtained the reaction of bird communities to urban load: communities of parks 6, 7, 9, and 10 constitute one similarity group (Fig. 5c). Thus, bird communities in forest parks, in which the urban recreation load is equal or higher than the median value, are combined into the same similarity group.

**Figure 5.**
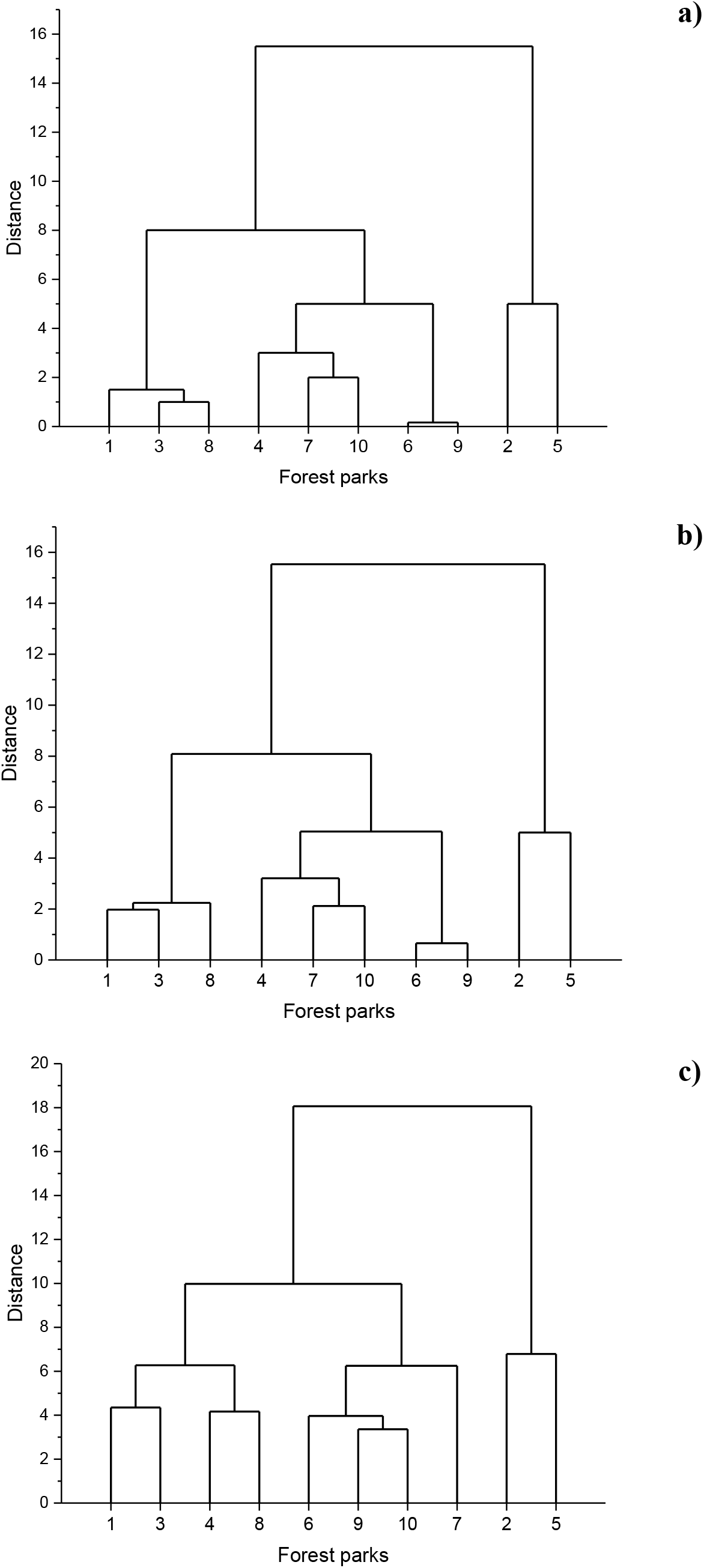
The similarity of nesting bird communities in Forest parks of the Kyiv: a) the result of the analysis of the α-diversity of communities; b) the result of analysis of nesting strategy; c) the result of a joint analysis of α-diversity indices and nesting strategy; see the sequence of forest parks in Figure 1

To identify the dependence of certain characteristics of nesting bird communities on the urban recreation load and the size of the forest park area, Principal Component Analysis was carried out. For the analysis, we took the characteristics of the bird community, the logarithmic trends of which were the most significant: number of tree canopy nesting species, undergrowth nesters, ground nesters, hollow nesters, synanthropy index, Shannon and Berger-Parker indices, the relative abundance of hollow nesters and ground nesters.

According to the results of PCA (Fig. 6), a positive correlation between the urban recreation load and the synanthropy index, the Berger-Parker index, the relative abundance of hollow nesters, and a negative correlation with the abundance of ground nesters have been revealed. The correlation between the urban recreation load and the synanthropy index of bird communities was the tightest.

**Figure 6.**
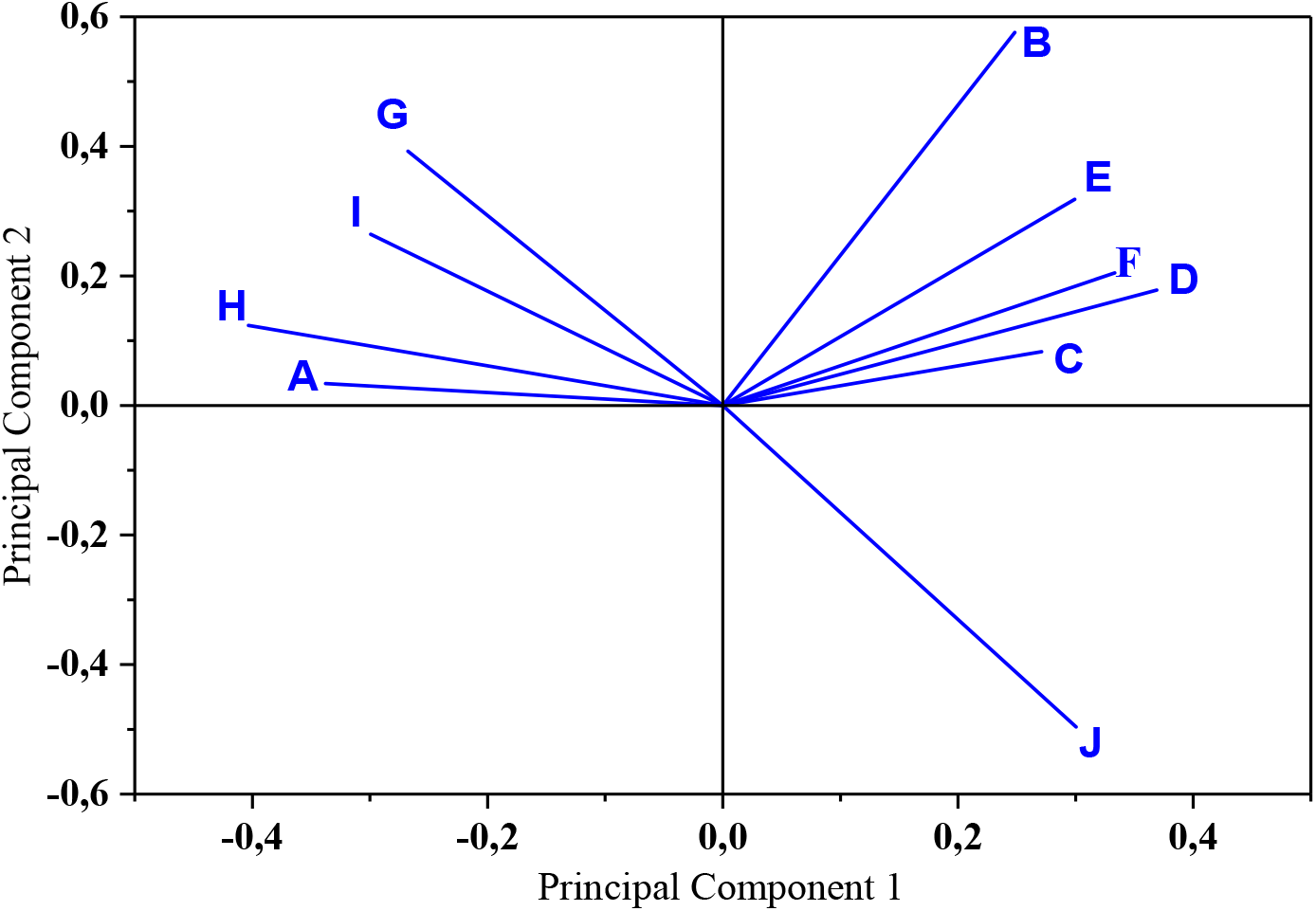
Principal Component Analysis: A – urban recreation load, B – number of canopy-nesting species, C – number of undergrowth-nesting species, D – number of ground-nesting species, E – number of hollow-nesting species, F – Shannon diversity index, G – Berger–Parker dominance index, H – the synanthropy index, I – the relative abundance of hollow nesters, J – the relative abundance of ground nesters

We also found strong associations between the Shannon diversity index and the number of the ground-nesting and undergrowth-nesting bird species.

## Discussion

In natural forests, the highest bird density was observed in structurally complex habitats with understorey and clutter (Felton et al., 2016; Blinkova and Shupova, 2018). In the transformed habitat, there is a shortage of safe nesting sites. Many species adapt to existence amid overpopulation in safe areas. Logarithmic trends in the number of nesting bird species of all 4 types of nesting strategies show a decrease along the line of increasing an urban recreation impact. This indicates that urban conditions have a detrimental effect on all forest bird species. It leads to a decrease in the number of nesting bird species, which will result in the development of communities with a small number of species and loss of species diversity in ecosystems of urban tree stands.

J.P. Hoover has emphasized interactions among food limitation, density, and reproduction (Hoover, 2020). Our research has shown a moderate value of the average density of birds nesting in most forest parks. Fluctuations in the average birds’ density are mainly associated with a significant difference in the dominant species density. It is worth paying attention to the differences in the nesting density of Blackbirds, which is widespread and dominant in most forest parks. Its nesting density in Kyiv forest parks is 2.5–14.5 pairs/ha. The importance of the dense bushes as forage stations of the Blackbirdis emphasized (Camprodon and Brotons, 2006; Orban et al., 2019). The synanthropic subpopulation of Blackbirds, brought from Poznan and acclimatized in 1958, settled sedentary in Kyiv (Gracek et al., 1975). We believe that settled individuals of synanthropic subpopulation and migratory individuals of non-synanthropic one inhabit forest sites on forest parks. Urbanization has facilitated the growth of the Blackbird population (Møller et al., 2014). Another dominant species in the forest parks of Kyiv is the chaffinch, the nesting density range of which is also wide: 3.2–25.5 pairs/ha. Chaffinch and Wood Pigeon (*Columba palumbus* L.) (0.7–3.0 pairs/ha) has synanthropized so much in Kyiv. The situation with these two species is interesting since in Catalonia Wood Pigeon and Chaffinch do not use forests with weak shrubbage (Camprodon and Brotons, 2006). At the same time, Wood Pigeon across much of Europe formed stable synanthropic subpopulations in the late XIX – early XX centuries (Tomiałojc, 1976), nesting in parks, squares, tree stands of streets.

An assessment of the correlation between bird communities and the structure of forest phytocoenoses of some of the analyzed forest parks (Blinkova and Shupova, 2018) depending on the recreational load intensity showed that the transformation of the forest area is only one element of the influence on the bird community evolution, in which the influence of surrounding biotopes associated with bird mobility is also important. J. Jokimäki (1999) described the dependence of the introduction of nesting species into parks from the landscape structure outside the park. The suitability of a forest is related to the surrounding landscape, and the response of birds to its fragmentation is not the same (Pretelli et al., 2018). The role of the surrounding landscapes in the development of bird communities has been also confirmed by this study. We have revealed for the first time for Kyiv that dendrophilous obligate synanthropes Serin (*Serinus serinus* Pallas) and Syrian Woodpecker (*Dendrocopos syriacus* Hemprich et Ehrenberg) settle on the outskirts of some forest areas, coming here from private house buildings or an amusement park bordering forest parks. Syrian Woodpecker and Serin were not previously observed in forests and forest parks. These birds are alien to the fauna of Ukraine.

We observed that forest-steppe species such as Gold Finch (*Carduelis carduelis* L.), Yellow Hammer (*Emberiza citrinella* L.), Common Linnet (*Linaria cannabina* L.) are typical for biotopes with ecotone effect began to nest in the forest parks of Kyiv. The phenomenon of homogenization of the species composition of avifauna in settlements has been described for cities of different regions (Chace & Walsh, 2006; Croci et al., 2008; Clavero and Brotons, 2010). V. Devictor et al. found out that landscape disturbance and fragmentation lead to functional homogenization of avifauna (Devictor et al., 2007). F. Morelli et al. have shown that urbanization leads to the disappearance of stenotopic species therefore urbanization planning is necessary (Morelli et al., 2016) to preserve habitat for all species (Vasileios et al., 2019).

In the forest parks of Kiev, some undergrowth nesters, change nesting strategy. Black bird, Eurasian Black Cap (*Sylvia atricapilla* L.), Greenfinch (*Chloris chloris* L.), Hawfinch (*Coccothraustes coccothraustes* L.) are more likely to nest at a height of more than 10 m in the crowns of the main forest-forming species, and only part of the population lives in the understorey (Gaychenko and Shupova, 2019). This analysis with a larger sample of forest parks confirmed the initial conclusions made in the work of V. Gaychenko and T. Shupova (2019). Thus, we have noted the change by part of the species its nesting strategy, and the redistribution of nesting gilds used by birds towards safer ones. A higher nest location of Blackbird has been described as a trait characteristic of urban populations (Luniak and Mulsow, 1988). We have shown that the negative effect of the urban recreation load becomes obvious to the good of birds using more protected nesting gilds, increasing the abundance of hollows nesters.

The negative relationship between urban recreation load and the relative abundance of ground nesters, revealed by Principal Component Analysis, has also confirmed previous studies. The urban recreation load was most tightly correlated with the synanthropy index of nesting bird communities, slightly less with the abundance of hollow nesters. R. Tryjanowski et al showed a positive correlation between the abundance of bird species and the area of tree stands as well as the age of trees. It has also been revealed that bird species richness was positively correlated with the area and age of trees (Tryjanowski et al., 2017). For India, using PCA, it was revealed that extension of the size of green spaces, an increase in the diversity and density of shrubs increases the number of forest bird species in cities (Khera et al., 2009). PCA revealed a positive correlation between the Shannon diversity index and the number of ground-nesting bird species as well as in the undergrowth-nesting ones in the forest parks of Kyiv.

The results of the cluster analysis have shown that the use of the characteristics of the α-diversity of communities and the nesting strategy of birds are equivalent in identifying similarity groups of communities and evolutionary connections within them. To identify the response of bird communities to urban recreation load, the best result is obtained by analyzing the complex of given α-diversity indices and characteristics of nesting strategy.

To preserve birds nesting on the ground and in the lower tiers of the stand, small species, birds that are not ready for synanthropization, as well as maintaining the species diversity of forest park communities, it is important to create forest areas protected from both human and animal influence. The steps for this are as follows: dense planting of shrubs and herbaceous plants, ornamental fences in forest areas with nesting birds that are at risk of accidental destruction of their nests. Fences should be accompanied by educational environmental advertisements on the importance of biodiversity conservation.

## Conclusion

Urban recreation load leads to a decrease in the number of nesting bird species of all types of nesting strategies due to the elimination of stenotopic representatives of forests. Forest-steppe, synanthropic, and alien species penetrate communities. The effect of urban recreation load contributes to a change in the nesting strategy of open nesters towards an increase in the height of the nest location. There is an increase in the relative abundance of cavity nesters, a decrease in the proportion of ground nesters, and the synanthropization of communities increases. To reveal the response of bird communities to urban recreation load, the best result is obtained by using a larger number of parameters (especially the complex of α-diversity indices and characteristics of ecological groups. According to the Principal Component Analysis, the strongest positive correlation of the urban recreation load with the synanthropy index, and a negative one with the abundance of ground nesters have been observed. To preserve vulnerable bird species (nesting on the ground and in the lower tiers of the stand, birds that are not ready for synanthropization) and maintaining the species diversity of communities, it is advisable to create forest areas in forest parks with densely planted shrubs and herbaceous plants so that they are almost impassable for humans and pets. It is also effective to put ornamental barriers prohibiting passage in areas of the forest with nesting birds that are at risk of accidental destruction of nests. Fences should be accompanied by environmental advertising on the need to preserve animals and their habitats.

## Acknowledgement

The authors express heartfelt appreciation to the doctor Olena Blinkova, for advice and the interpretation and presentation of Principal Component Analysis results and PhD. Sergej Koniakin, a research fellow of the Institute for Evolutionary Ecology of the NAS of Ukraine for help in drawing up a Kyiv map.

